# A Strategy for Differential Abundance Analysis of Sparse Microbiome Data with Group-wise Structured Zeros

**DOI:** 10.1101/2023.07.24.549296

**Authors:** Fentaw Abegaz, Davar Abedini, Fred White, Alessandra Guerrieri, Anouk Zancarini, Lemeng Dong, Johan A. Westerhuis, Fred van Eeuwijk, Harro Bouwmeester, Age K. Smilde

## Abstract

Comparing the abundance of microbial communities between different groups or obtained under different experimental conditions using count sequence data is a challenging task due to various issues such as inflated zero counts, overdispersion, and non-normality. Several methods and procedures based on counts, their transformation and compositionality have been proposed in the literature to detect differentially abundant species in datasets containing hundreds to thousands of microbial species. Despite efforts to address the the large numbers of zeros present in microbiome datasets, even after careful data preprocessing, the performance of existing methods is impaired by the presence of inflated zero counts and group-wise structured zeros (i.e., all zero counts in a group). We propose and validate using extensive simulations an approach combining two differential abundance testing methods, namely DESeq2-ZINBWaVE and DESeq2, to address the issues of zero-inflation and group-wise structured zeros, respectively. This combined approach was subsequently successfully applied to two plant microbiome datasets that revealed a number of taxa as interesting candidates for further experimental validation.

## Introduction

The plant and soil microbiomes, comprising a diverse community of beneficial and harmful microbes, play an important role in plant growth and health^1–3^. To understand the mechanisms that govern plant-microbiome interactions, high-throughput sequencing methods have been dramatically advanced^4^. Amplicon sequencing (e.g., 16S rRNA gene) and whole genome shotgun sequencing are the two major methods for the identification of microbial communities by utilizing sequence data^5, 6^. The resulting microbiome data consisting of counts and compositional data, are typically characterized by sparsity (zero-inflation), overdispersion (variance that is much higher than expected), high-dimensionality (number of taxa is much higher than the number of samples), non-normality, and variable sequencing depth among samples. Such characteristics of the microbiome data make its analysis challenging^7^. However, failure to consider these special characteristics of microbiome count data during statistical analysis can result in false positive results and irreproducible relationships^8^.

One important aspect of microbiome data analysis that attracts significant scientific interest is the detection of differential abundance among microbial species across two or more conditions or treatments. However, microbiomes consist of hundreds to thousands of distinct taxa (a term used to refer to OTUs: operational taxonomic units or ASVs: amplicon sequencing variants), with just a small percentage likely to be differentially abundant^9^. Several methods and procedures based on counts, compositionality, and transformation perspectives have been introduced in the literature for discovering differentially abundant species (see^10, 11^ and references therein for a list of methods used in differential abundance analysis). Current practices for detecting differential abundance in microbiome data involve careful data pre-processing (filtering and normalization) and the use of suitable statistical tools that consider the special characteristics of the data^12, 13^. However, there is a continued debate over the appropriate approaches for assessing differential abundance in microbiome data^12^.

Despite strict quality control and contaminant removal utilizing QIIME^14^ and DADA2^15^ software, microbiome data still contain many rare and low-prevalence taxa and are thus highly zero-inflated^16^, which makes microbiome data analysis challenging. In microbiome data, typically between 80 and 95 percent of the counts are zero^12^, while, the number of zeros varies significantly across taxa, ranging from none or a few to many zeros. Zero counts in the sample can simply reflect absence but also presence with low frequency that was not detected due to technical detection limits. In particular, zeros in sequence count data can be either biological zeros which indicate the true absence of taxa under specific environmental conditions, or non-biological zeros which can arise from various factors such as sequencing errors, limited sequencing depth, uneven sampling depth, and PCR amplification bias^14, 17–20^. Unfortunately, without prior biological knowledge or spike-in controls, distinguishing biological and non-biological zeros in sequence count data is difficult^17^. In spite of this, reviews cited in^10^ demonstrated that many rare taxa are a result of sequencing artifact contamination, and/or sequencing errors^14^ are not informative in the analysis and have no influence on the scientific conclusions. The inclusion of a filtering step to remove potentially uninformative taxa before performing statistical tests can reduce the burden of adjusting for multiple tests, which considerably improves detecting differentially abundant taxa^11^. Several ad hoc and principled filtering algorithms have been presented in this respect. The choice of filtering strategy will influence the results of the subsequent analysis^12^.

While filtering reduces the complexity of microbiome data, the highly-inflated-zeros that remain even after filtering can result in significant reductions in statistical power if not adequately modeled^15^. Several statistical tools have been introduced to properly deal with the analysis of zero-inflation, including zero-inflated Gaussian^16^ and zero-inflated negative-binomial^21^ models, as well as observation weights-based strategies for using popular RNA-seq tools such as DESeq2, edgeR, and limma-voom^22^. The inflated-zeros, on the other hand, directly contribute to another statistical issue, the problem of perfect separation^23^ or group-wise structured zeros^24^, which arises when many non-zero counts are in one group and all zero counts in the other. When there is enough evidence to believe that biological factors caused the occurrence of taxa with group-wise structured zeros they can be identified as structural zeros and labelled as significant without further differential abundance testing because they are abundant in one group but not at all present in the other group^24^. On the other hand, because it is difficult to identify biological from non-biological zeros in sequence count data, it would not be fair to relate group-wise structured zeros to only biological zeros. For instance, the presence of group-wise structured zeros is more likely to be noticeable in small samples but it is possible that group-wise structured zeros may not be present in the population as a whole or with more sampled data. In such circumstances, the occurrence of group-wise structured zeros needs be explored to see if it is caused by an inherent biological factor or sampling variability. To this end, implementing standard likelihood inference in the presence of taxa with perfect separation or group-wise structured zeros, hampers statistical inference. This is because it results in large or infinite parameter estimates of effects coupled with extremely inflated standard errors, causing such taxa to be nonsignificant^23^.

In the statistical literature, a penalized likelihood strategy that provides finite parameter estimates has been suggested as a solution to the issues that perfect separation or group-wise structured zeros present when using maximum likelihood-based techniques^23, 25^. Furthermore, penalized likelihood ratio-based tests help in providing appropriate significance test results for perfectly separated taxa^23, 25^. However, no or little efforts have been made to include penalized likelihood inference into many of the existing differential abundance techniques. Among the differential abundance techniques, the estimation approach implemented in DESeq2 by combining ridge type penalized likelihood estimation with a likelihood ratio based test^26^ has the potential to address the problem of perfect separation or group-wise structured zeros.

Differential abundance analysis is generally conducted using a single or combination of methods adopted from bulk RNA sequencing analysis, single-cell RNA sequencing analysis, or particularly developed for microbiome data^16^. There have been a few large-scale benchmarking studies that look at the adequacy of using these methods^10, 11, 27^. According to the benchmarking studies, the various differential abundance analysis methods are inadequate at controlling false discovery rates at nominal levels, and there was no consistency in the opinion what the right approach is^21^. Here we focus mainly on popular tools that take into consideration the count nature of the data, such as edgeR^28^, DESeq2^26^, limma-voom^29^, as well as their weighted counterparts referred to as edgeR-ZINBWaVE, DESeq2-ZINBWaVE and limma-voom-ZINBWaVE^22^, which utilize a weighting mechanism based on the ZINBWaVE model^30^. Some of these and other approaches were evaluated on a vast variety of microbiome datasets^10^. However, many of these techniques treat taxa with group-wise structured zeros differently in differential abundance testing. To ensure a fair comparison of these techniques, taxa with group-wise structured zeros were identified and excluded from the differential abundance testing in our simulated comparisons.

In this work, first, we considered a simulation strategy similar to Mallick et al.^8^ that mimics experimental plant microbiome data using the simulation model SparseDOSSA^6^ with several performance metrics to compare weighted and unweighted differential abundance methods; second, we implemented a combination of differential abundance tools that include (i) DESeq2-ZINBWaVE: ZINBWaVE-weighted methods to address the problem of zero inflation and control false discovery rate and (ii) DESeq2: penalized likelihood ratio based method to properly address the analysis of taxa with perfect separation or group-wise structured zeros; third, we created a comprehensive pipeline for differential abundance analysis of microbiome data that includes data pre-processing and identification of differentially abundant taxa using the combined approach DESeq2-ZINBWaVE and DESeq2; finally, we applied the pipeline for detecting differential abundance on two experimentally obtained plant microbiome 16S rRNA gene sequencing datasets from the MiCRop (Microbial Imprinting for Crop Resilience; www.microp.org) project.

## Methods and materials

### Pipeline design

The combined approach, DESeq2-ZINBWaVE-DESeq2, is designed to perform a thorough assessment of microbial abundance differences while accounting for zero-inflation and group-wise structured zeros. Fig 1 depicts the essential stages in the implementation of DESeq2-ZINBWaVE-DESeq2. The input data includes an abundance table, a taxonomy table, and a metadata table in any standard microbiome data format. The initial stages of data processing involve filtering and normalization. For the analysis of a single treatment with two factor levels, the data pre-processing step is readily followed by categorizing taxa as having group-wise structured zeros or not. Here we note that if there is sufficient evidence to assume that taxa with group-wise structured zeros are due to biological causes, they can be labelled as significant without further differential abundance testing^24^. Otherwise, we will proceed as follows. For each category, differential abundance analysis is done independently (Analysis Part A and B). In Analysis Part A, DESeq2 likelihood ratio test (LRT) is used to perform differential abundance testing for taxa with group-wise structured zeros while in Analysis Part B, DESeq2-ZINBWaVE-based LRT is utilized for taxa without group-wise structured zeros. Finally, we collect significant taxa from both analyses for diagnostic purposes and biological interpretation. On the other hand, when the identification of group-wise structured zeros becomes more difficult, for example, in the presence of multiple categorical factors, the structural zero grouping step can be skipped. Without much loss of power, the entire filtered and normalized dataset can be analyzed using both DESeq2 (which is appropriate in detecting differential abundance for taxa with group-wise structured zeros) and DESeq2-ZINBWaVE (which is more powerful in detecting differential abundance for taxa with no group-wise structured zeros) based LRTs. This is followed by collecting unique significant taxa from both analyses for diagnostic purposes and biological interpretation.

**Figure 1:**
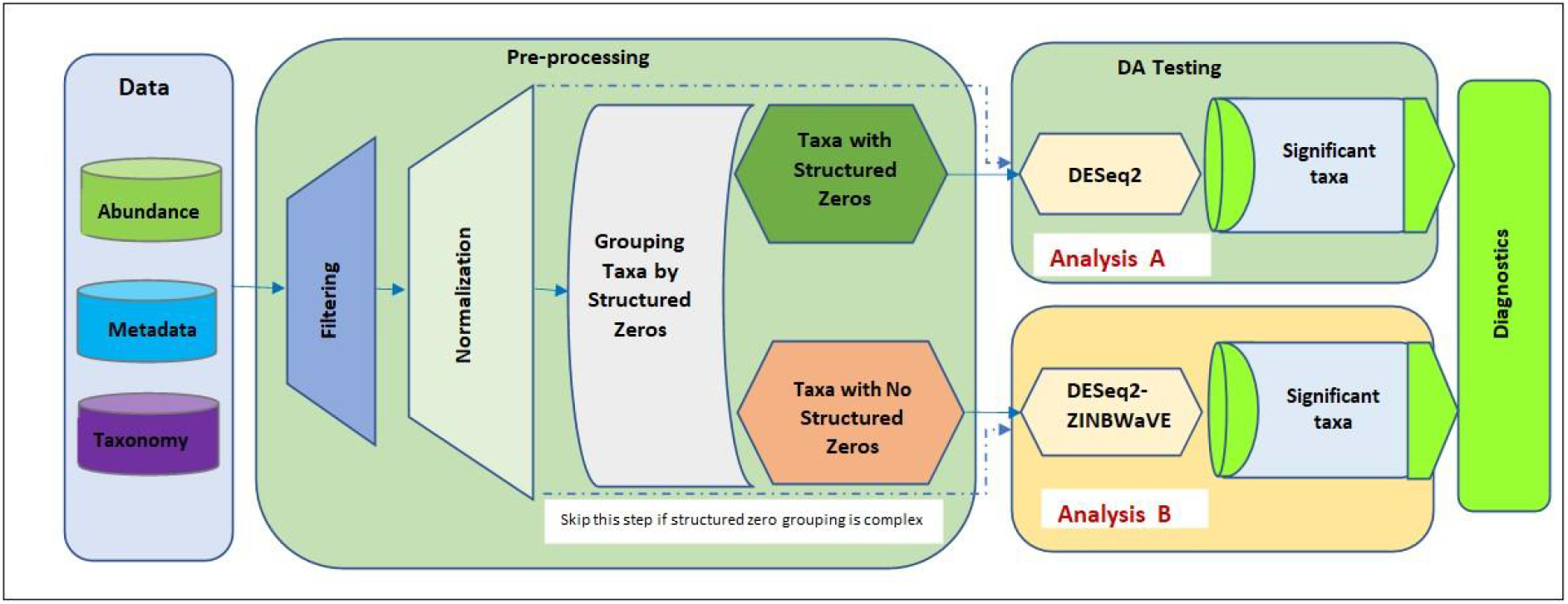
Flowchart for the Microbiome Data Differential Abundance Analysis Pipeline. **Data:** the input microbiome data include the abundance table, taxonomy table, and metadata table. **Pre-processing step**: includes filtering and normalization. Grouping taxa by group-wise structured zeros for a single covariate with two factor levels. **Differential abundance testing** can be performed in two parts. **Analysis Part A**: differential abundance testing for taxa with group-wise structured zeros using DESeq2 likelihood ratio test (LRT). **Analysis Part B**: differential abundance testing for taxa without group-wise structured zeros using DESeq2-ZINBWaVE with LRT. **Diagnostics**: collect significant taxa from both analyses for diagnostic purposes and biological interpretation. When there are multiple covariates, skip the structural zero grouping step and apply both DESeq2 and DESeq2-ZINBWaVE on the whole filtered and normalized data (dashed arrows).

#### Data preparation

Following raw sequence read processing with DADA2 or QIIME, the sequencing data is typically presented in two table formats: an abundance table for counts and a taxonomy table for phylogenetic information. Differential abundance analysis generally requires the use of three input datasets: (i) Abundance data table: taxa count abundance across samples; (ii) Taxonomy data table: taxonomy information across taxa (required for higher hierarchical levels of analysis or interpretation of results); and (iii) Metadata table: information on treatments, phenotypes or covariates of interest across samples (Fig. 1). It is important to note that, as part of the data preparation step, we must examine the abundance and metadata tables for missing data values. We considered two template datasets differing in the level of zero-inflation for evaluating the performance of popular unweighted and weighted microbiome differential abundance tools (see details in the data analysis section).

#### Data Filtering

Because microbiome data sets are usually sparse, it is necessary to filter the data set by removing low-quality or uninformative taxa to improve downstream statistical analysis. Several filtering approaches are implemented in R/Bioconductor packages that include filter_taxa in phyloseq^31^, filterByExpr in edgeR^28^, and simultaneous or permutation based statistical filtering in PERFect^12^. The edgeR filterByExpr function allows filtering based on replications or treatment groups. Unless indicated, our filtering to retain a taxon for differential abundance analysis is based on a minimum of two counts in at least two samples per treatment group.

#### Taxa Grouping

Following the filtering stage, taxa are grouped based on whether they have group-wise structured zeros or not. Taxa with group-wise structured zeros have zero counts in all samples of one of the groups.

#### Data Normalization

Normalization is another important step in microbiome sequencing data analysis that is used to remove any bias caused by differences in sequencing depths or library sizes between samples. For microbiome studies, two forms of normalization have been used: rarefying and scaling^9, 32^. Rarefying is subsampling to equal sequencing depth without replacement^9^. We did not use rarefying-based normalization since its application in differential abundance analysis is debatable^11, 33^. Scaling *based* normalization: this is to acquire a scaling factor that can be used to adjust the read counts to produce normalized counts or to produce normalized library sizes^34^. Normalized library sizes are used as offsets in count-based regression models such as DESeq2 and edgeR and their weighted counterparts to remove biases caused by uneven sequencing depths in differential abundance analysis ^34^. Some commonly used scaling-based normalization procedures include Cumulative-Sum Scaling (CSS) implemented in metagenomeSeq, Relative Log Expression (RLE) in DESeq2, Upper Quartile (UQ) in limma-voom, and Trimmed Mean of M-values (TMM) in edgeR. These procedures were developed primarily for RNA-Seq data that suffer less of large numbers of zeros than microbiome data. To address zero-inflation in normalization, geometric mean of pairwise ratios (GMPR)^34^, geometric mean of positive counts (poscounts) and deconvolution ^35^ methods were introduced; however, normalization methods for zero-inflated microbiome data are still under development. In our analysis, we implemented the commonly used TMM and poscounts normalization for edgeR/limma-voom and DESeq2, respectively. In our simulation studies, we observed that GMPR normalizations demonstrated comparable performance to poscounts in controlling false discovery rate using DESeq2 and DESeq2-ZINBWaVE (Fig S2).

#### Data exploratory analysis

##### Evaluating the extent of zero inflation

one of the primary challenges in microbiome differential abundance analyses is the presence of excessively inflated zero counts^14^. We used a graphical depiction of the biological coefficient of variation^22^, which is the square root of the estimated negative binomial dispersion parameter, to examine the extent of inflated-zeros and how they influence downstream differential abundance analyses. Taxa with few counts or many zeros result in very high dispersion estimates, which appear as striped patterns on biological coefficient of variation plot. The increased dispersion due to inflated zeros hampered the capacity to detect differential abundance using negative binomial based models such as edgeR and DESeq2^22^, which do not account for excess zeros.

#### Type I error control

Another way to assess the suitability of a differential abundance method is to evaluate its type I error rate control using mock or model-based simulated data. On the one hand, we generated mock samples from real null data that did not include taxa whose abundance differed between two groups (i.e., no differentially abundant taxa between two groups), by randomly assigning the two groups to each taxon. On the other hand, we generated 100 datasets using the SparseDOSSA model with an effect size of zero, i.e., under the null hypothesis of no differentially abundant taxa between two groups. Filtering, grouping by group-wise structured zeros, and normalization were applied to the mock and simulated datasets. Taxa containing group-wise structured zeros were removed from differential abundance testing in the simulation studies. Then, using several methods, we conducted a differential abundance analysis between two groups of the covariate and obtained the p-values. The proportion of the p-values less than the commonly used nominal 5% level is used to calculate the observed type I error rates. For each differential abundance approach, the average observed type I error rates across mock or model-based simulated datasets were computed.

#### Differential abundance testing

Many methods for analyzing differential abundance in microbiome data are adapted from methods developed for analyzing bulk RNA-Seq or single cell RNA-Seq data^10^. Differential abundance approaches tailored to the study of microbiome data have also been developed. In our comparative study, we looked at a few common approaches for detecting differential abundance that have previously been tested using multiple real microbiome datasets^10, 11, 36^ and synthetic abundance data^6^. We considered microbiome differential abundance analysis approaches adapted from RNA-Seq analysis techniques, such as edgeR, DESeq2, and limma-voom as well as their weighted counterparts: DESeq2-Zimbwave, edgeR-ZINBWaVE, and limma-voom-ZINBWaVE, respectively, recently introduced for single-cell RNA-Seq, which employ weights generated using the ZINBWaVE model^10, 22^. We also included ANCOM-BC from the compositional approach and MaAsLin2 and metagenomeSeq from transformation-based microbiome data analysis approaches.

#### Plant microbiome experimental datasets

In this work, we used plant microbiome datasets with varying degrees of zero-inflation and count pattern obtained from two experimental studies to assess several differential abundance techniques.

#### Plant material and DNA extraction

In the N-P starvation experiment, the impact of nitrogen (N) and phosphate (P) starvation on the bacterial composition in tomato (*Solanum Iycopersicum*) roots was investigated. *S. lycopersicum cv.* Moneymaker (SS687) seeds were surface sterilized using 70% EtOH (2 min), 20% bleach (20 min), 10mM HCl (10 min) and milli-Q water (5 times), respectively. The seeds were then pregerminated in a climate room at 24°C in the dark on a sterilized and moisturized filter paper for 3 days. The germinated seeds were transferred to soil-filled baskets in the greenhouse at 22°C under an 8/16 h dark/light regime. Ten days after germination, the baskets were transferred to a custom-made aeroponics system, in which the roots were sprayed with 1/4 strength Hoagland solution for 15 seconds every 10-minutes. For the N starvation treatment, NH4NO3 was removed from the solution. For the P starvation treatment, K2HPO4 was replaced with 0.8M KCL, which compensated for the loss of K. The roots were collected and immediately transferred to liquid nitrogen and stored at −80°C for further analysis.

In the Forest-Potting soil experiment, the effect of different soil types on the bacterial community of tomato roots was investigated. To this end, tomato seeds were surface-sterilized as described above and then sown in different soils with or without the addition of 10% forest soil. After four weeks, the roots were collected and used for microbial DNA extraction.

#### Bacterial DNA isolation, 16S rDNA sequencing and preprocessing

The frozen root tissues were ground into a fine powder using liquid nitrogen and a mortar and a pestle. 200 mg of the powdered tissue was used to extract microbial DNA using PowerSoil DNA Isolation following the manufacturer’s instructions. The quality and quantity of the isolated DNA were assessed using a Qubit 2·0 Fluorimeter (ds DNA high-sensitivity assays kit; Invitrogen). The metabarcoding analysis was conducted at BaseClear B.V. (Leiden, The Netherlands), utilizing an Illumina NovaSeq6000 SP platform with the 2×250 bp paired-end sequencing approach for samples from the N-P experiment. For the amplification of bacterial 16S rDNA, the V3 and V4 regions were targeted using the 341F (5’- CNTACGGGNGGCWGCAG-3’) and 805R (5’- GACTACHVGGGTWTCTAATCC-3’) primers. The sequencing reads were obtained as demultiplexed reads and after quality check, the primers were trimmed from the reads using cutadapt (v.1.9.1). For samples from the Forest_Potting soil experiment, the metabarcoding analysis was conducted on Illumina MiSeq PE250 platform at Génome Québec Innovation Centre (Montréal, QC, Canada). The sequencing data were processed using DADA2 package and the plugin Quantitative Insights Into Microbial Ecology version 2 (QIIME2) to generate an OTU count table. The reads were filtered to maintain 220 bp and 200 bp for the forward and reverse reads, respectively. After merging denoised pair-end sequences, the chimeric sequences were removed from the reads. The obtained Operational Taxonomic Units (OTUs) were taxonomically assigned based on the SILVA database (v138), and an OTU count table was created for each dataset.

#### Datasets

The N-P starvation dataset included 2945 taxa with at least one count from 5 samples in each of the three treatment groups: N-starvation, P-starvation and control. We filtered out low abundance taxa using the criteria of at least two counts in at least two samples per treatment group, which resulted in 829 taxa for differential abundance analysis. The percentage of zero counts was around 55% after filtering. The Forest-Potting soils dataset included a total of 28 samples with 1796 taxa that had at least one count from each of the two soil types: Forest and Potting soils. The Forest soil type includes different soils with the addition of at least 10% forest soil. We filtered out low abundance taxa, leaving 244 with a minimum of two counts in at least two samples of each soil type for differential abundance analysis. Even after filtering, the fraction of zero counts remained high (84%), indicating that the Forest-Potting soils dataset is considerably zero-inflated as compared to the N-P starvation dataset. Moreover, very low taxa counts in the Forest-Potting soils dataset were another feature that distinguishes it from the N-P starvation dataset.

### Simulation studies

#### Model-based simulation studies

To evaluate the performance of our method and others on differential abundance detection for microbiome data, we used the SparseDOSSA model to generate synthetic data^6^. This model has the benefit of generating realistic simulated data based on parameterizing real-world template microbial datasets targeting the main characteristics of microbiome data such as zero-inflation, counts, compositionality, and high dimensionality^6^.

##### Simulating null synthetic taxa abundance data

In the model-based simulation setting, we generated null abundance taxa data (no significant differential abundance taxa related to a treatment or covariate of interest) with varying total sample sizes (10, 20, 50, 100, 200) that mimic the structure of Forest-Potting soils and N-P starvation real data templates from plant microbial community using sparseDOSSA2^6^. The generated null synthetic datasets were used to evaluate the performance of differential abundance methods in controlling type I error rate.

##### Simulating synthetic metadata

For synthetic metadata generation, we considered a single factor with two levels. The metadata was generated by randomly assigning a value of 1 to half of the samples and a value of 0 to the other half.

##### Simulating spike-in taxa to describe synthetic taxa-metadata relationships

To introduce differences in abundance between two groups, varying effect sizes or log-fold changes (0.5, 1, or 2) were used. The following is how we produced synthetic data with spike-in taxa based on real template data. First, we selected about 10% of taxa in the template data that had a relative abundance of at least 20%. Then, for these taxa (called spike-in taxa), treatment effects were assigned of known effect sizes or log-fold changes: 0.5, 1 or 2. Finally, the sparseDOSSA2 R/Bioconductor package was used to generate simulated count data for the spike-in taxa according to their specified effect sizes.

#### Mock dataset based studies

For the mock studies, we generated 1000 datasets with two groups for both the Forest-Potting soils dataset (potting and forest soil) and the N-P starvation dataset (N-starvation and control) with 14 and 5 samples each, respectively.

#### Performance evaluation

Mainly two performance indicators are used for evaluation: statistical power or sensitivity and False Discovery Rate (FDR), which are computed based on false positives (FPs): taxa not spiked-in but found significant, true positives (TPs): taxa spiked-in and found significant, true negatives (TNs): taxa not spiked-in and not found significant), and false negatives (FNs): taxa spiked-in but not found significant.

#### Pipeline implementation

The pipeline (Fig. 1) provides a comprehensive differential abundance analysis of microbiome data, including data preparation, filtering, normalization, differential abundance testing and diagnostic plots. The entire pipeline is written in R. Several R and Bioconductor packages are used in the pipeline to analyze and visualize differential abundance detection using DESeq2, edgeR, limma-VOOM, and their ZINBWaVE weighted counterparts: DESeq2-Zimbwave, edgeR-ZINBWaVE, and limma-voom-ZINBWaVE.

In the pipeline, following the test of significance of all filtered taxa for differential abundance using one of the methods listed; log-fold changes, p-values, and adjusted p-values that are corrected for multiple hypothesis testing using Benjamini-Hochberg are provided in a summary table. Plots depicting statistically significant differential abundant taxa are also generated. A variety of summary and diagnostic plots are also provided to visualize significant results in the pipeline: (i) plots of significant taxa vs. log fold change; (ii) plots of log fold change vs. average log CPM (counts per million) for all taxa; (iii) count plots to evaluate significant taxa; and (iv) heatmap plots for significant taxa with relative abundances.

## Results

### Zero-inflation and perfect separation or group-wise structured zeros

#### Popular differential abundance tools handle perfect separation or group-wise structured zeros differently

In order to investigate how group-wise structured zeros are handled and how they impact differential abundance detection, we analyzed the N-P starvation plant microbiome dataset using DESeq2 and ZINBWaVE-weighted DESeq2. Following data filtering, poscounts normalization and identification of taxa with group-wise structured zeros based on observed zero counts (see the methods and materials section), first we performed differential abundance testing using DESeq2 and DESeq2-ZINBWaVE on the entire set of taxa. We observed a substantial variation in the number and type of significantly differentially abundant taxa identified by DESeq2 and DESeq2-ZINBWaVE (Figure 2). Figure 2, highlights taxa depending on their statistical significance and whether or not they are involved in perfect separation or group-wise structured zeros. Results from DESeq2 (Fig. 2A) show that many taxa with group-wise structured zeros were found significant with substantial log-fold changes by lying on the boundary of the plot (cyan dots in Fig. 2A). Using DESeq2-ZINBWaVE which down-weights excess zeros, in contrast, no or a few taxa with group-wise structured zeros were found to be significant for the N-P starvation data as displayed in Fig. 2B. This is not necessarily the case using DESeq2-ZINBWaVE, since we found many taxa with group-wise structured zeros to be significant after reanalyzing the Arctic-soil data, which contains a large number of samples investigated in [8] (Fig. S1B).

**Figure 2:**
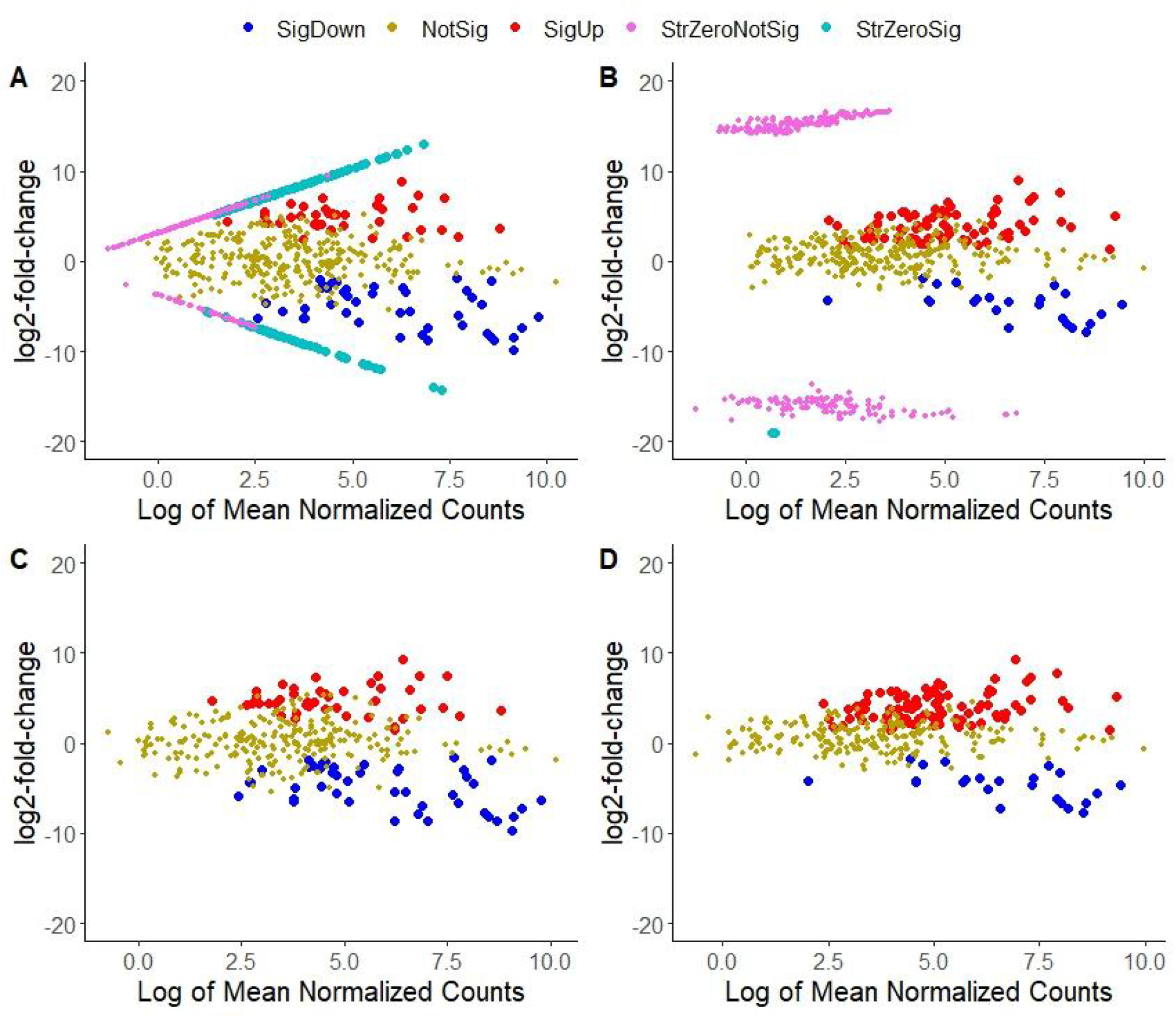
Comparing differential abundance detection tools in the presence of perfect separation or group-wise structured zeros for the N-P starvation dataset comparing Nitrogen starvation to control. SigDown: significant taxa with negative log-fold change, SigUp: significant with a positive log-fold change, NotSig: not significant, StrZeroSig: significant for taxa with group-wise structured zeros, StrZeroNotSig: not significant for taxa with group-wise structured zeros. A. Analysis with DESeq2, taxa with group-wise structured zeros found to be significant having relatively large log-fold changes and located on the boundary of the plot (cyan); B. Analysis with DESeq2-ZINBWaVE, taxa with group-wise structured zeros found not to be significant (purple) because of down weighting excess zeros. The number of significant taxa identified by DESeq2 and DESeq2-ZINBWaVE differed considerably due to the presence of taxa with group-wise structured zeros. C. Analysis with DESeq2 after excluding taxa with group-wise structured zeros; D. Analysis with DESeq2-ZINBWaVE after excluding taxa with group-wise structured zeros.

Because there was a large difference in the number of significant taxa, which was mostly attributed to taxa with group-wise structured zeros in this case, we reanalyzed the data using DESeq2 and DESeq2-ZINBWaVE after excluding taxa with group-wise structured zeros. The findings shown in Fig. 2 (C and D) demonstrate that DESeq2 and DESeq2-ZINBWaVE detect a comparable number of taxa (red and blue dots), which may occur when zero-inflation is not a serious concern ^22^. As a result, comparative benchmarking studies on microbiome differential abundance analysis tools that do not account for group-wise structured zeros could lead to unsatisfactory comparative conclusions.

#### Type I error control

We assessed the type I error rate using mock samples and model-based synthetic data derived from the two real template datasets under the null hypothesis of no differentially abundant taxa between two groups. (see the methods and materials section).

In the model-based simulation with SparseDOSSA2 under the null hypothesis, we used a log-fold change of zero, implying that no taxa were differentially abundant between the two groups. We then performed differential abundance analysis and recorded the p-values for several methods under consideration. The observed type I error rates were calculated based on the fraction of p-values smaller than the nominal 5% level. The resulting plots shown in Fig 3 for model based simulations and Fig S2 for the mock samples, reveal how each method controls type I error under the null hypothesis of no differentially abundant taxa. Simulation results based on the N-P starvation template dataset with moderately inflated-zeros (Fig 3A), DESeq2, edgeR, MaAslin2, DESeq2-ZINBWaVE (for moderate and large samples), edgeR-ZINBWaVE (for small samples) and limma-voom (for small samples) demonstrated effective control of the type I error rate at the nominal level. DESeq2-ZINBWaVE (for small samples) and edgeR-ZINBWaVE (for large samples), on the other hand, had slightly higher observed type I error rates. In contrast, simulation findings using the Forest-Potting soils template dataset with highly inflated-zeros (Fig 3B) demonstrated that DeSeq2-ZINBWaVE, edgeR-ZINBWaVE and MaAsLin2 controlled type I error rate at the nominal level. On the other hand, DESeq2 and edgeR were conservative tests with extremely small observed type I error rates, which might impede true discoveries. In comparison to the other methods, the observed type I error rate for limma-voom-ZINBWaVE was much higher, showing poor performance in terms of controlling the type I error rate on both synthetic datasets. The performance of DESeq2-ZINBWaVE (for small samples) and edgeR-ZINBWaVE (for large samples) in controlling type I error rate improved on average as zero-inflation was increased.

**Figure 3.**
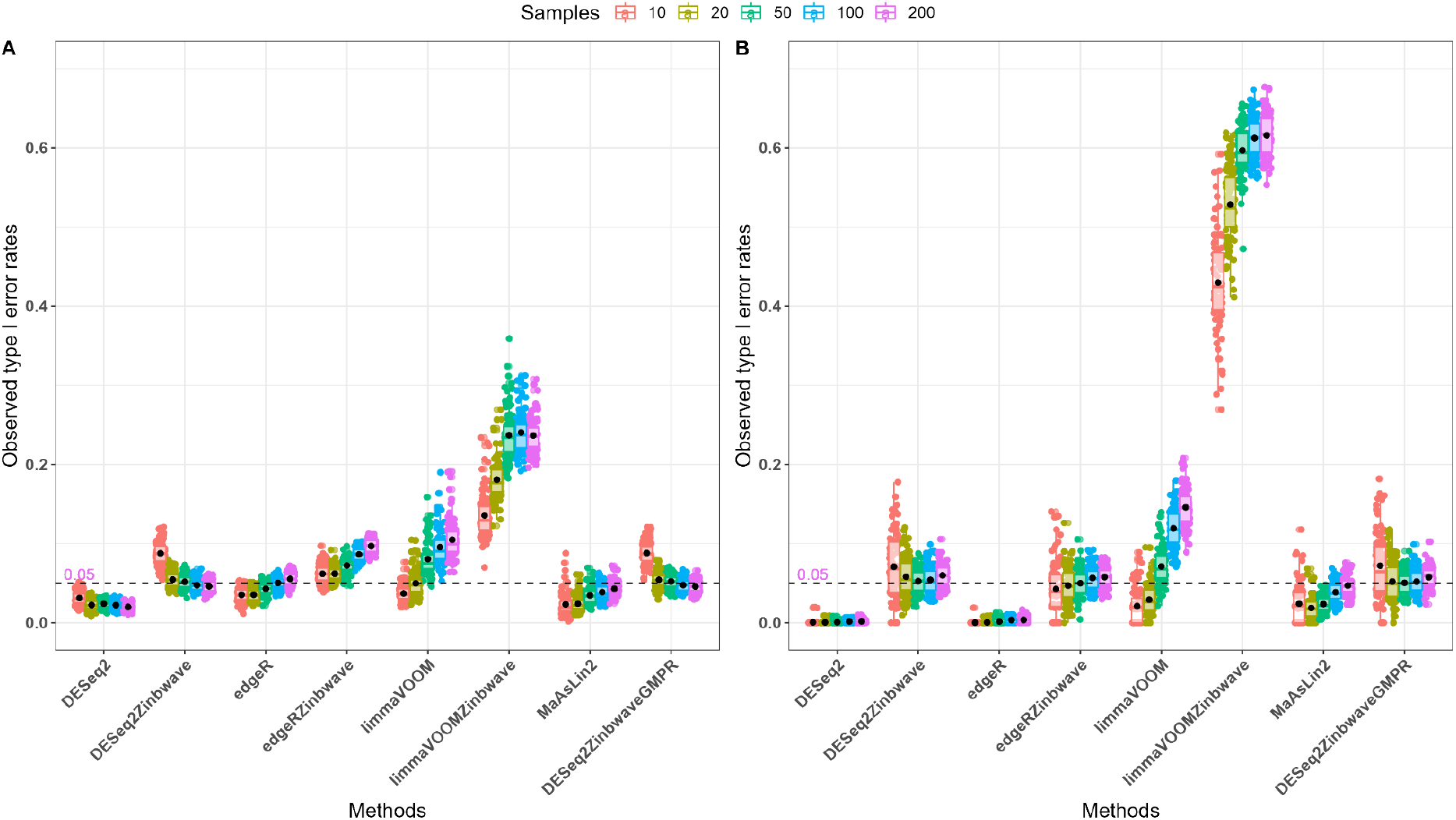
Model-based simulations: controlling type I error rates using several differential abundance tools. Unweighted and weighted differential abundance methods were evaluated for type I error control based on synthetic plant microbiome data with varying zero-inflation rates and sample sizes. Boxplots of observed type I error rates are colored by total sample size. A: N-P starvation template dataset with 55% zeros. B: Forest-Potting soils template dataset with 84% zeros. Weighted approaches to differential abundance, with the exception of limma-voom-ZINBWaVE, demonstrated acceptable control type I error rates for moderately and highly zero-inflated datasets.

Similar results were observed based on 1000 mock datasets generated from N-P starvation and Forest-Potting soils datasets (Fig S2). On the other hand, the observed type I error rates for limma-voom-ZINBWaVE, ANCOM-BC, and metagenomeSeq were high on both mock datasets, potentially leading to a large number of false discoveries (Fig S2 A and B). As a result ANCOM-BC and metagenomeSeq were not included in the model-based simulation results presented above. Moreover, using GMPR instead of poscounts and TMM normalizations had comparable impact on controlling type I error rate.

#### Differential abundance methods benchmarking using synthetic data

We utilized synthetic count data with spiked-in taxa between two defined groups of samples to evaluate the performance of differential abundance analysis tools (see the methods and materials section). We simulated 100 datasets, each with a single treatment with two factor levels and a fixed number of true differential features selected based on at least 20% of differentially abundant taxa for varying effect sizes (0.5, 1.0, 2.0) and total sample sizes (10, 20, 50, 100, 200). The biological coefficient of variation plots computed from the template real data and simulated data are shown in Fig. S3 to demonstrate how the data generating process resembles the structure of the N-P starvation template real dataset.

We then used false discovery rate (FDR) and power (sensitivity) as performance indicators to assess the ability of different microbiome differential abundance analysis approaches to recover the relationship between spike-in taxa and a two-factor treatment. In the simulation analysis, following filtering, normalization and removing taxa with group-wise structured zeros (to place methods in comparable context), tests of differential abundance were performed for each of the 100 datasets using each of the methods under consideration. Based on the adjusted p-values (Benjamini-Hochberg), we identified true positives (TP), false positives (FP), true negatives (TN) and false negatives (FN) and computed the measures of performance such as power and FDR. For each method, the power and FDR are displayed in Fig. 4 and 5.

**Figure 4.**
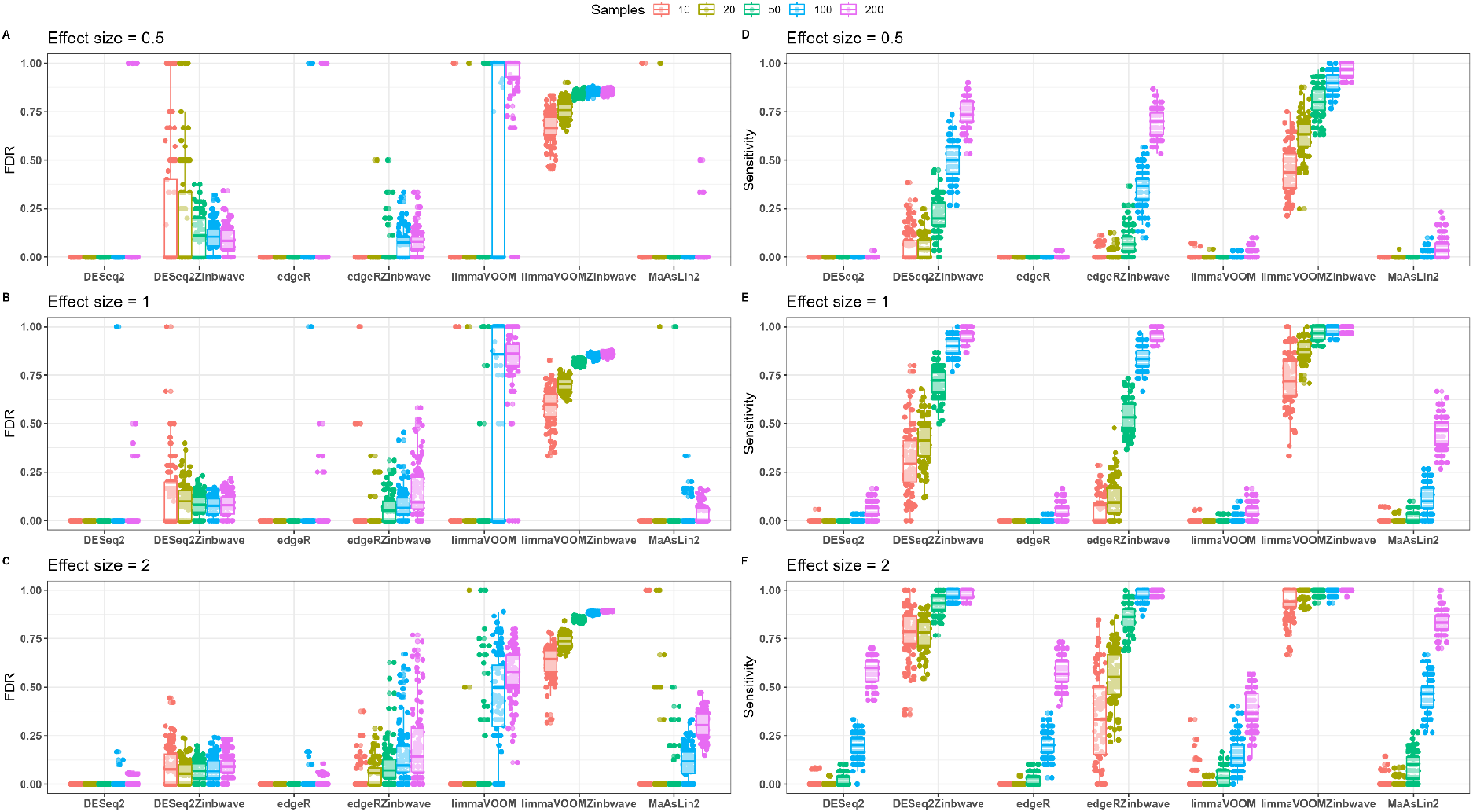
FDR and statistical power (sensitivity) for weighted and unweighted differential abundance detection methods evaluated on a highly zero-inflated Forest-Potting soils template plant microbiome data. Boxplots of FDR (A, B, C) and power (D, E, F) with effect sizes of 0.5 (A, D), 1 (B, E) and 2 (C, F) are colored by total sample size. DESeq2-ZINBWaVE demonstrated a FDR close to the nominal 5% level the nominal level while maintaining power at reasonably high level except in a very small sample case. The power of limma-voom-ZINBWaVE was high but at the expense of a very high FDR. The performance of the unweighted methods DESeq2, edgeR and limma-voom was very poor for highly zero-inflated data.

**Figure 5.**
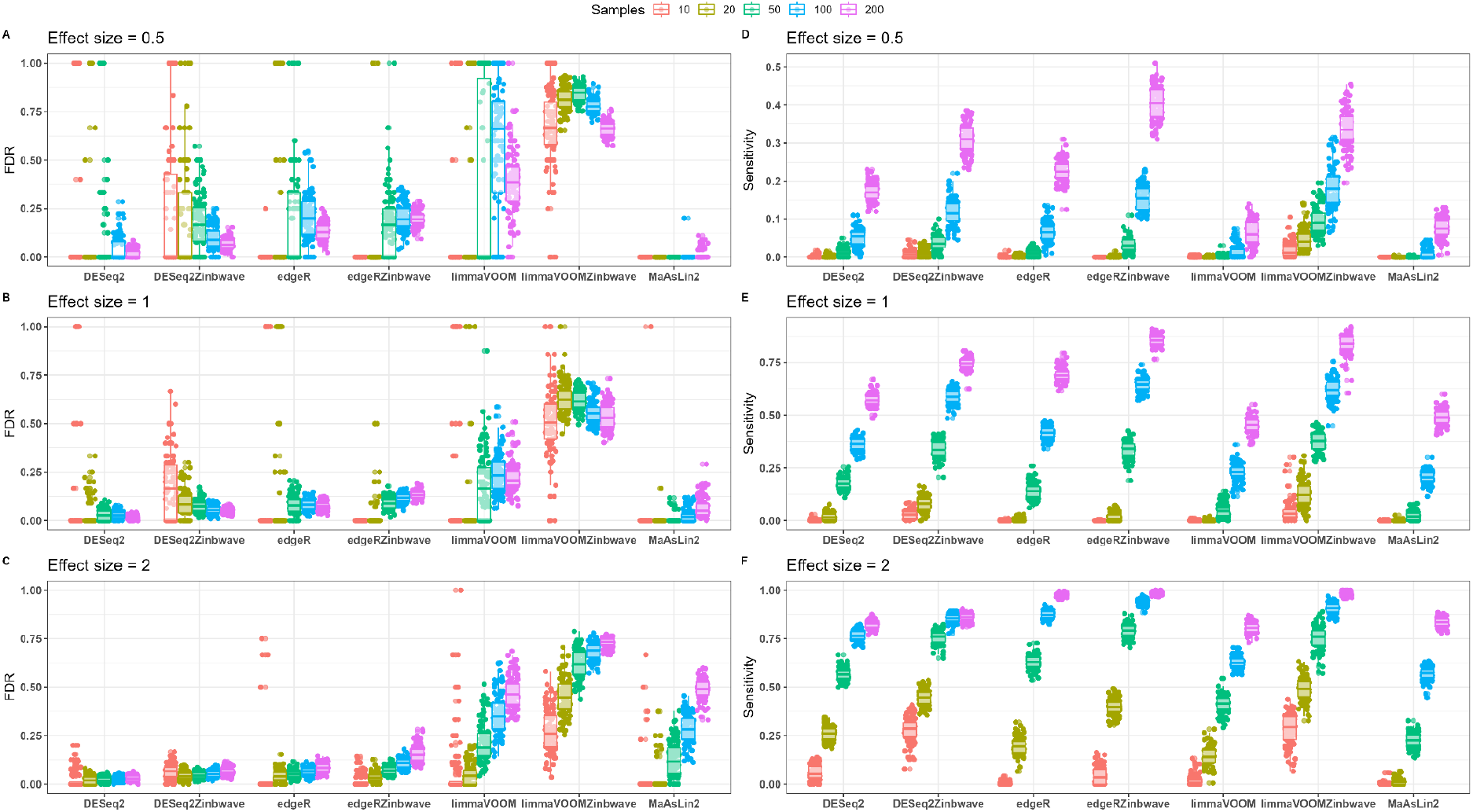
FDR and statistical power for weighted and unweighted differential abundance detection methods evaluated on moderately zero-inflated synthetic N-P starvation plant microbiome data. Boxplots of FDR (A, B, C) and power (D, E, F) with effect sizes of 0.5 (A, D), 1 (B, E) and 2 (C, F) are colored by total sample size. Unweighted methods demonstrated comparable performance with their weighted counterparts.The FDR of unweighted methods was on average slightly lower than their weighted counterparts but this was at a slight expense on their power. The comparable power of limma-voom and limma-voomZINBWaVE was overshadowed by a very high FDR.

##### Weighted methods outperform in highly zero-inflated data

For the highly zero-inflated Forest-Potting soils synthetic datasets, the simulation performance of differential abundance approaches using FDR and statistical power is shown in Fig 4. Except for limma-voom-ZINBWaVE, DESeq2-ZINBWaVE and edgeR-ZINBWaVE all methods evaluated had very poor statistical power (a very few or no true positives found significant) for small sample sizes across all effect sizes. Moreover, even with large sample and effect sizes, DESeq2, edgeR and limma-voom exhibited low power. In general, the power of all methods increased with increasing sample and effect sizes. The FDR values of DESeq2 and edgeR were mainly either 0 (no false positives were found) or 1 (all significant results were false positives). DESeq2-ZINBWaVE, on the other hand, resulted FDRs that were close to the nominal 5% on average except for small samples and small effect sizes. However, the FDR of DESeq2-ZINBWaVE decreased with increasing sample and effect sizes. For modest sample sizes, edgeR-ZINBWaVE demonstrated FDR values on average close to the nominal level, with a tendency to rise with sample size. Limma-voom-ZINBWaVE, on the other hand, demonstrated high power even in small sample sizes. However, in all sample and effect size settings, the FDR of limma-voom-ZINBWaVE exceeds the nominal threshold by a wide margin. That is, with a nominal FDR of 5%, on average, more than 60% of the features identified as significant by limma-voom-ZINBWaVE were false positives (Fig. 4). Across all samples and effect sizes, DESeq2-ZINBWaVE outperformed the other approaches in terms of power while maintaining a reasonable FDR control.

##### Better power of weighted methods even in moderately zero-inflated data

We also examined the FDR controlling and power behavior of different differential abundance detection methods for moderately zero-inflated N-P starvation synthetic datasets. Except limma-voom, limma-voom-ZINBWaVE and MaAslin2, the FDR in all other methods were close to the nominal 5% level (Fig. 5) for large effect and sample sizes. The FDR of unweighted methods was on average slightly lower than their weighted counterparts but this was at a slight expense on power. DESeq2-ZINBWaVE showed a relatively better power performance than edgeR-ZINBWaVE in small sample sizes. For large samples, on the other hand, edgeR-ZINBWaVE had better power than DESeq2-ZINBWaVE but this was at the expense of increased FDR.

##### The potential of utilizing DESeq2-ZINBWaVE-DESeq2 approach for differential abundance analysis

To assess the performance of the combined approach DESeq2-ZINBWaVE-DESeq2, we reanalyzed the metagenome shotgun sequencing data from the Human Microbiome Project (HMP-2012), which included 5 supragingival and 5 subgingival plaque samples from the oral cavity^10, 37^ (see details in Text S1). Using enrichment analysis, DESeq2-ZINBWaVE-DESeq2 discovered many accurate enrichments when compared to the other differential abundance tools (Fig. S5).

#### Analysis of experimental datasets

##### Forest-Potting soils microbiome analysis

Out of 244 filtered taxa occurring in at least two replicates from one of the two soil types and with a minimum of two counts per taxon, 135 had group-wise structured zeros and 109 did not. DESeq2-ZINBWaVE-DESeq2 was applied to detect differentially abundant taxa between forest and potting soils. We used DESeq2 on 135 taxa that had group-wise structured zeros and DESeq2-ZINBWaVE on 109 taxa that did not have group-wise structured zeros. Fig. 6 and Fig. S4 depict the taxa identified as differentially abundant between potting and forest soils (adjusted p-value < 0.05). In Fig 6, the relative abundances are colored, with higher relative abundances represented by dark red. The figures show that mainly specific *Massilia* and *Streptomyces* ASVs were more abundant in potting and forest soils, respectively.

**Fig 6.**
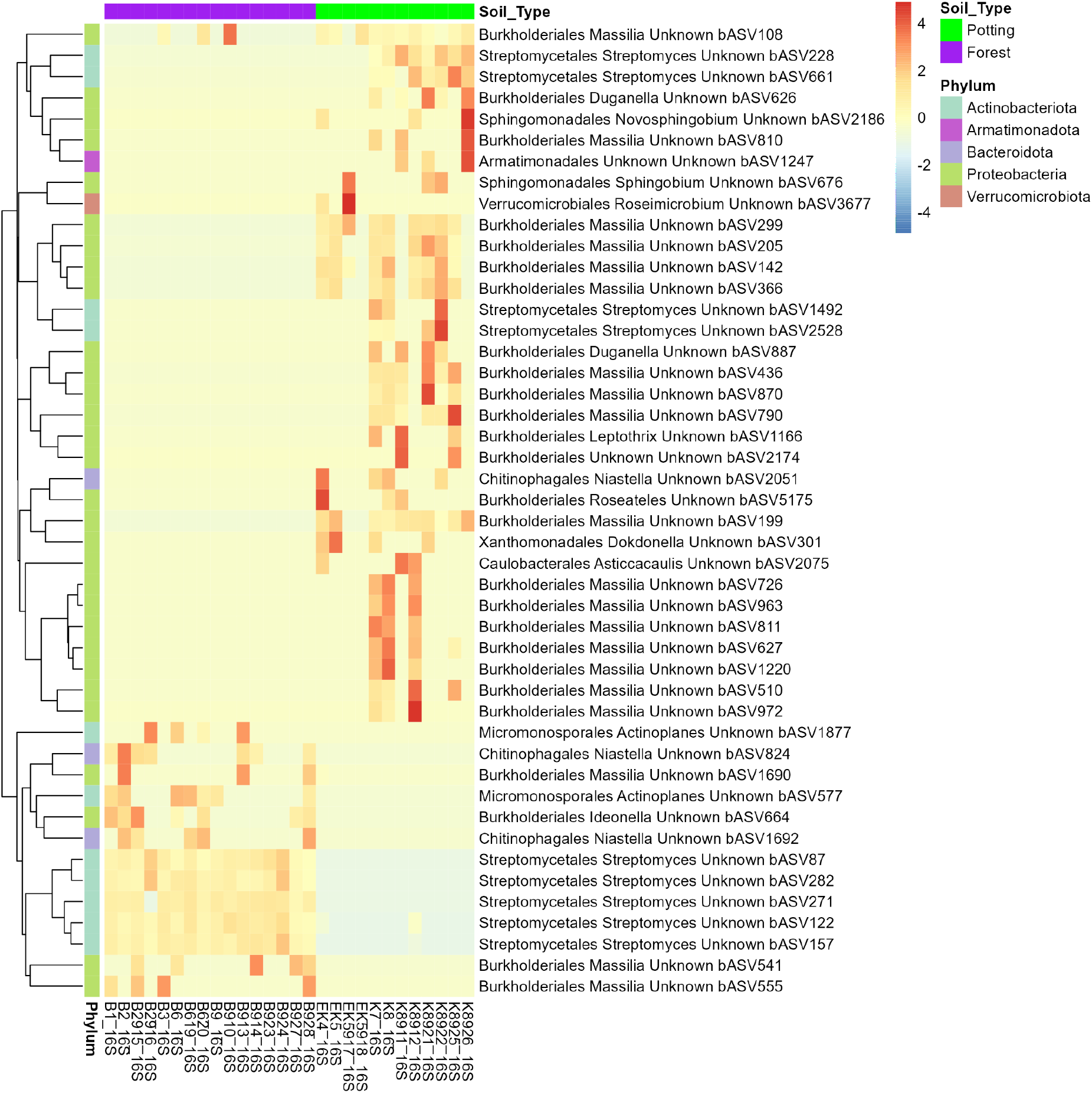
Heatmap of differentially abundant taxa between potting and forest soils. Higher relative abundances represented by darker red.

##### N-P starvation microbiome analysis

We also applied the combined DESeq2-ZINBWaVE-DESeq2 approach to detect differentially abundant microbes in tomato growing under nitrogen (N) and phosphate (P) deficiency compared to a control with complete Hoagland solution (C). To utilize the likelihood ratio test in DESeq2 and DESeq2-ZINBWaVE for multicategorical variables, we modified the design matrix so that the generated dummy variables could be easily included into the full and reduced model structures, comparable to the likelihood ratio implementation in edgeR-ZINBWaVE^30^. As a result, the full model structure contains the intercept, N versus C, and P versus C, but the reduced model can include the intercept and either N versus C or P versus C. In comparing N versus C, we had 451 taxa with group-wise structured zeros and 320 without. Similarly, for P versus C, there were 385 taxa with group-wise structured zeros and 444 without. Fig. 7 depicts the taxa identified as differentially abundant comparing N versus C and Fig. 8 comparing P versus C using the methods DESeq2 for taxa with group-wise structured zeros and DESeq2-ZINBWaVE for taxa without group-wise structured zeros (adjusted pvalue < 0.05). Under N-starvation taxa belonging to the families: *Mycobacteriaceae, Caulobacteraceae, Comamonadaceae, Bdellovibrionaceae, Rhizobiaceae, Streptomycetaceae, Reyranellaceae, Hyphomonadaceae, Candidatus_Kaiserbacteria* and *Candidatus_Nomurabacteria* were more abundant while taxa belonging to the families: *Acetobacteraceae, Acidobacteriaceae, Microbacteriaceae, Burkholderiaceae*and *Rhodanobacteraceae* were less abundant. Under P-starvation taxa belonging to the families: *Sphingomonadaceae, Acetobacteriaceae, Bdellovibrionaceae, Caulobacteraceae* and *Candidatus_Kaiserbacteria* were more abundant while taxa belonging to the families: *Microbacteriaceae, Intrasporangiaceae, Xanthobacteracea* and *Rhizobiales* were less abundant.

**Figure 7.**
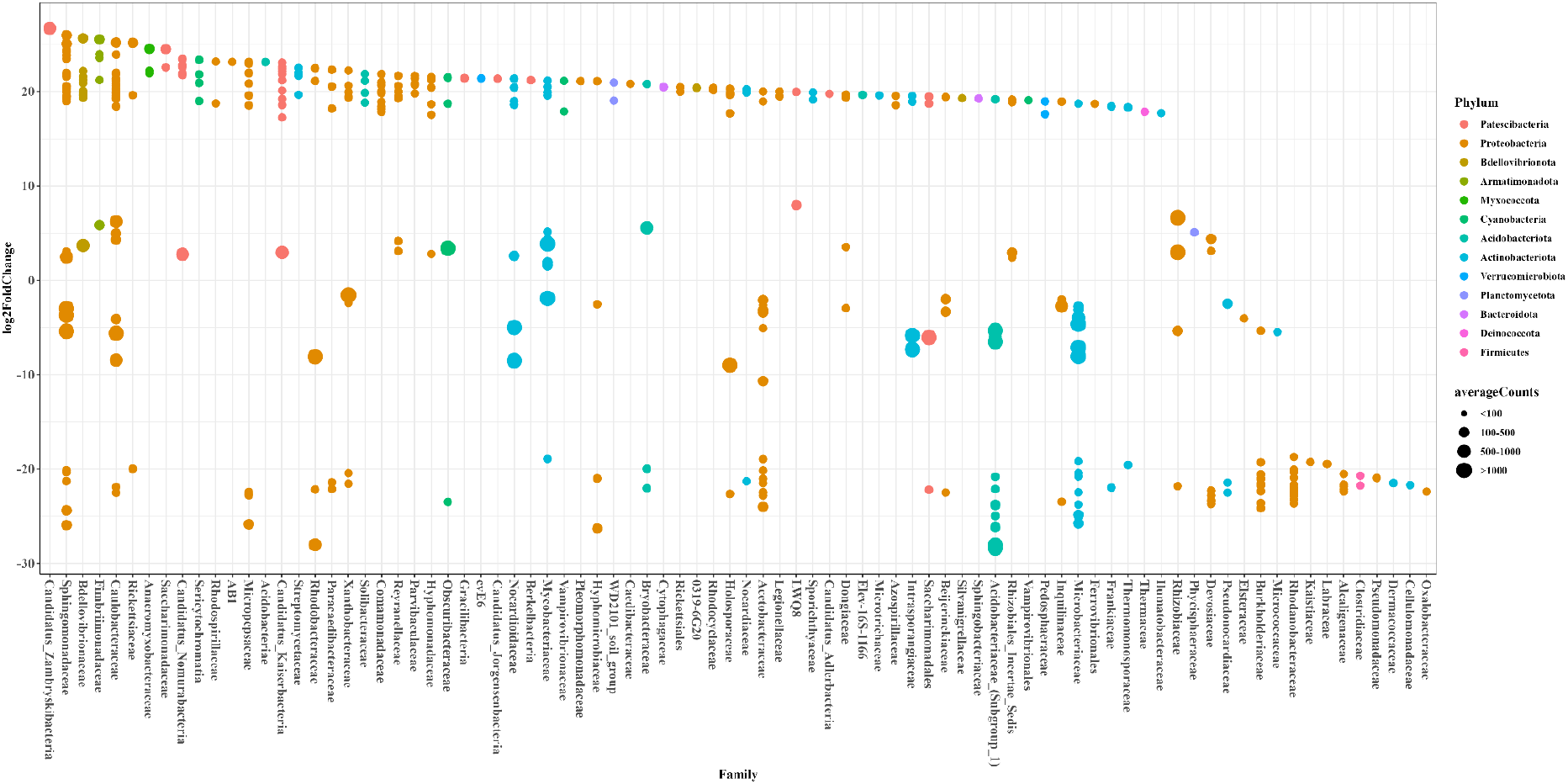
Differentially abundant taxa in comparing N-starvation with the control.

**Figure 8.**
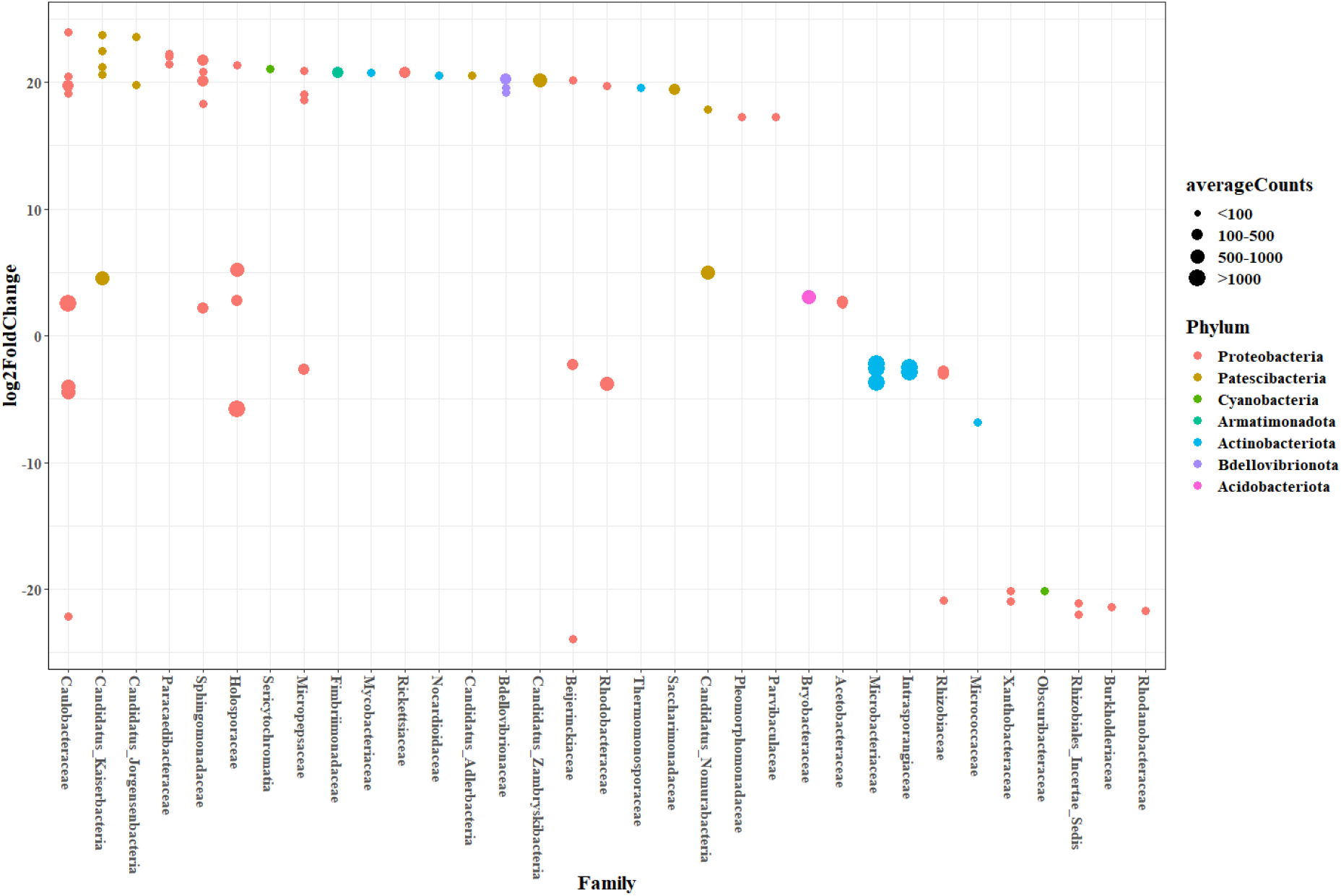
Differentially abundant taxa in comparing P-starvation with the control.

## Discussion and conclusion

This study aimed to contribute to the search for the best count-based differential abundance practices for microbiome data. In microbiome research, experimental datasets usually only have small to moderate number of replicates or total sample size. Moreover, in addition to inflated zeros, the microbiome count data may contain many taxa with group-wise structured zeros. The causes of group-wise structured zeros might be biological or non-biological factors, but identifying them in sequence count data is difficult. As a result, applying differential abundance methods in the presence of taxa with group-wise structured zeros makes statistical inference problematic due to extremely inflated standard errors, leaving many taxa nonsignificant. We included a pre-processing step to distinguish taxa with and without group-wise structured zeros as much as possible and analyze them separately. We implemented the DESeq2-ZINBWaVE-DESeq2 approach, which is a combination of differential abundance tools that include DESeq2-ZINBWaVE for analyzing taxa without group-wise structured zeros and DESeq2 for analyzing taxa with group-wise structured zeros.

With the zero-inflated microbiome data, we found a considerable difference in the number and type of significantly differentially abundant taxa using unweighted (DESeq2, edgeR, limma-voom-ZINBWaVE) and weighted (DESeq2-ZINBWaVE, edgeR-ZINBWaVE, limma-voom) differential abundance analysis techniques. In this regard, the handling of taxa with group-wise structured zeros in the various tools had a significant influence. The likelihood ratio-based tests in the unweighted methods produced many significant taxa with group-wise structured zeros, but the weighted counterparts produced none or a few, which may be attributed to down-weighting the excess zeros that might have distorted the contribution of zeros to perfect separation. In either the weighted or unweighted approaches considered, no or little attention was given to directly address the issue of group-wise structured zeros. As a result, comparative benchmarking studies on microbiome differential abundance analysis tools that do not account for group-wise structured zeros may yield incorrect results.

We next investigated the impact of zero-inflation on the power and FDR control performance of weighted and unweighted approaches in microbiome differential abundance analysis using mock samples and model-based synthetic datasets. Instead of being dependent on the underlying distributional assumptions of the methodologies under consideration, the model-based simulation experiments were designed to mimic real microbiome data using the SparseDOSSA model. We used two experimental plant microbiome datasets with moderately and highly inflated zeros. Noting the inconsistencies in how the various tools handled taxa with group-wise structured zeros (all zero counts in one of the groups) and to create a common platform for comparison, in the simulation analysis, taxa with group-wise structured zeros were excluded from the differential abundance testing. In our simulation study, we mainly compared frequently used differential abundance methods such as DESeq2, edgeR, and limma-voom to their ZINBWaVE weighted counterparts, DESeq2-ZINBWaVE, edgeR-ZINBWaVE, and limma-voom-ZINBWaVE^10, 22^. ANCOM-BC, metagenomeSeq, and MaAsLin2 were also examined to some extent. We investigated the finite-sample properties of these methods, focusing on their performance on false discovery and detection power. A range of sample and effect sizes were taken into account.

According to our simulation assessment, ZINBWaVE weighted DESeq2 or edgeR demonstrated reasonable power of detecting differential abundance in substantially zero-inflated microbiome data with moderate to large sample and effect sizes. Thus, utilizing weights to downweight excess zeros by adapting the popular RNAseq methods is a useful strategy for analyzing microbiome data. However, like the other differential abundance tools, ZINBWaVE weighted approaches had low power in detecting spike-in taxa with small effects and small sample sizes. This highlights the need for conducting power analysis when planning microbiome investigations, which helps in determining sample sizes while keeping the required power in mind^8, 10^. Moreover, inaccurate estimates of weights might have a negative impact on differential abundance detection. Here we considered ZINBWaWE-based weighting, which uses a zero-inflated negative binomial model, but alternative weighting schemes could be considered.

On the other hand, consistent with findings in previous studies, (i) type I error control was satisfactory under the null hypothesis of no differential abundance using the mock samples as well as the model-based simulated datasets when inflated zeros were properly accounted for; (ii) the FDR in all the methods we considered to identify spiked-in taxa were on average higher than the nominal level in the analysis of the zero-inflated microbiome data^8^. In this regard, while our simulation findings showed that employing weighted techniques to discover differential abundance is a step forward, controlling FDR remains a challenge in microbiome data analysis, necessitating continued refining of existing methods or the development of new ones.

Furthermore, the simulation experiments demonstrated that increasing sample size increased FDR in some of the differential abundance analysis methods evaluated. In particular, FDR control of edgeR-based methods were not improved by increasing sample size combined with large effect sizes. This could be a result of the bias involved in the estimated effects, which increases with sample size^24^. This end, the implementation of methods based on penalized likelihood inference in model parameter estimation could help to alleviate the problem of inflated FDR control.

Another possible solution for reducing the inflated FDR would be proper exploitation of the hierarchical nature of microbiome data. Recent findings in the literature offer several strategies for leveraging hierarchical structure to boost the identification of differentially abundant species. In this regard, methods were introduced using smoothing p-values according to the phylogeny^13^ or correlation tree^5^ and using hierarchically adjusted p-values^5, 38, 39^. However, the inclusion of phylogenetic information in microbiome differential abundance analysis generated inconsistent results in terms of detection power and FDR control^5, 13^ necessitating the development of novel tools.

The performance of the combined approach was assessed using enrichment analysis. In comparison to the independent analyses performed by DESeq2-ZINBWaVE and edgeR-ZINBWaVE, the combined technique DESeq2-ZINBWaVE-DESeq2 discovered many correct enrichments.

Finally, DESeq2-ZINBWaVE-DESeq2 was applied to investigate the two plant microbiome datasets utilized as templates for the data simulation. Our new approach identified many potentially important taxa that might be further explored in terms of effect sizes and abundance in order to prioritize them for biological validation. In conclusion, the combined method DESeq2-ZINBWaVE-DESeq2 described in this study provides a promising development in the analysis of microbiome datasets displaying zero-inflation and group-wise structured zeros.

## Acknowledgements

We acknowledge funding by the Dutch Research Council (NWO/OCW) for the MiCRop Consortium program, Harnessing the second genome of plants (Grant number 024.004.014; to HB, LD, DA, AKS, JAW, FVE and FA), the Dutch Research Council (NWO-TTW grant 16873 Holland Innovative Potato; to HB, LD and DA), the ERC (Advanced grant CHEMCOMRHIZO, 670211; to HB, AZ and AG) and the Data Science Centre of the University of Amsterdam (to FW).

## Data availability

All data sets generated and analyzed in the current study can be available upon reasonable request to the authors. The R-code used to analyze the data will be available in GitHub repository.

## Author contributions

FA, AKS, JAW and FVE conceptualized the methodology and study design. FA implemented the method, performed the analysis and wrote the draft manuscript. AKS, JAW and FVE provided feedback throughout the manuscript preparation. DA, FW, AG, AZ, LD and HB designed the plant experiments; processed, sequenced and provided the microbiome data. All authors provided critical revisions and approved the final manuscript.

## Supplementary information

**S1 Fig. Comparing differential abundance detection tools in the presence of perfect separation or structural zeros for Arctic fire soil.** SigDown: significant taxa with negative log-fold change, SigUp: significant with a positive log-fold change, NotSig: not significant, StrZeroSig: significant for taxa with structural zeros, StrZeroNotSig: not significant for taxa with structural zeros. A. Analysis with DESeq2, taxa with structural zeros found to be significant having relatively large log-fold changes and located on the boundary of the plot (cyan); B. Analysis with DESeq2-ZINBWaVE, taxa with structural zeros found not to be significant (purple). The number of significant taxa identified by DESeq2 and DESeq2-ZINBWaVE differed considerably due to the presence of taxa with structural zeros. C. Analysis with DESeq2 after excluding taxa with structural zeros; D. Analysis with DESeq2-ZINBWaVE after excluding taxa with structural zeros.

**S2 Fig. Mock samples: controlling type I error rates using several differential abundance tools.** Unweighted and weighted differential abundance methods were evaluated for type I error control based on 1000 mock samples from plant microbiome data with varying zero-inflation rates. Left panel: N-P starvation template dataset with 55% zeros. Right panel: Forest-Potting soils template dataset with 84% zeros. Compared to the 5% nominal level, on average the observed type I error rates were very high for ANCOM-BC, metagenomeSeq, and limma-voom-ZINBWaVE; very low for DESeq2, edgeR and MaAsLin2; slightly higher for DESeq2-ZINBWaVE N-P starvation dataset; and close to for DESeq2-ZINBWaVE Forest-Potting soils dataset and for edgeR-ZINBWaVE and under the null hypothesis of no differentially abundant taxa.

**S3 Fig. Biological coefficient of variation (BCV) plots obtained from template real data and synthetic data**. The BCVs are displayed against the average number of log counts per million (AveLogCPM). The BCV plot displays striped patterns, which are a sign of taxa with many zeros and high estimates of dispersion. For the N-P starvation template real dataset, the BCV estimates from the template real data and one simulated data are comparable to each other.

**S4 Fig. Differentially abundant taxa in comparing potting and forest soils.**

**S5 Fig. Enrichment analysis comparing Supragingival vs Subgingival plaque for the metagenome shotgun sequencing samples from the Human Microbiome Project (HMP-2012). Each bar represents the number of significantly (adjusted p-value < 0.10) abundant taxa by each method, with positive log fold changes in Supragingival (top red bars), negative log fold changes in Subgingival plaque (bottom blue bars), and positive log fold changes in Supragingival (top blue bars), coloured according to aerobic and anaerobic metabolism. Fisher exact test is employed to establish the enrichment significance, and the p-values are shown.**

**S1 Text. The potential of utilizing the DESeq2-ZINBWaVE-DESeq2 approach for differential abundance analysis**

(pdf)

