## Supplemental information for "A Strategy for Differential Abundance Analysis of Sparse Microbiome Data with Group-wise Structured Zeros"

### Figures S1-S5 and Text S1

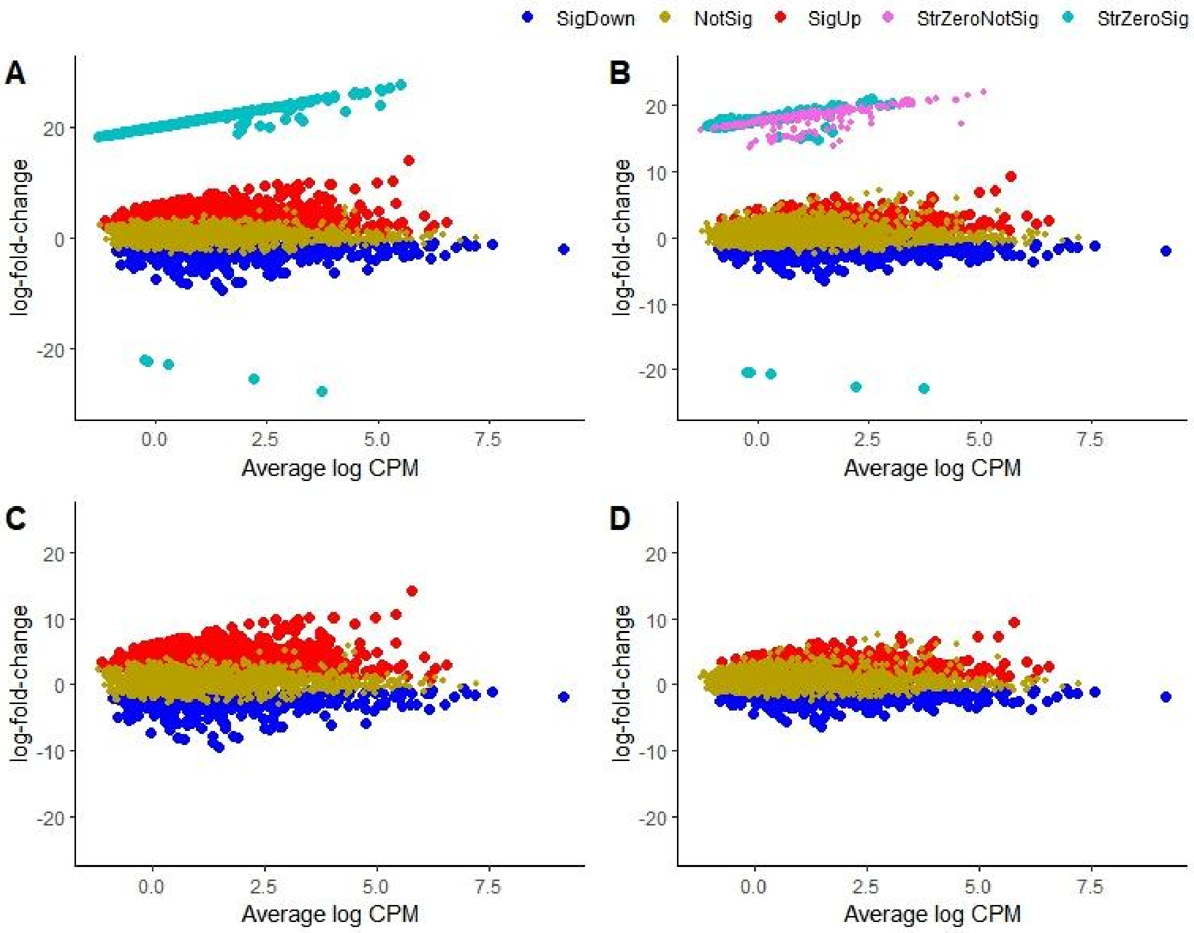

**Figure S1: Comparing differential abundance detection tools in the presence of perfect separation or structural zeros for Arctic fire soil.** SigDown: significant taxa with negative log-fold change, SigUp: significant with a positive log-fold change, NotSig: not significant, StrZeroSig: significant for taxa with structural zeros, StrZeroNotSig: not significant for taxa with structural zeros. A. Analysis with DESeq2, taxa with structural zeros found to be significant having relatively large log-fold changes and located on the boundary of the plot (cyan); B. Analysis with DESeq2-ZINBWaVE, taxa with structural zeros found not to be significant (purple). The number of significant taxa identified by DESeq2 and DESeq2-ZINBWaVE differed considerably due to the presence of taxa with structural zeros. C. Analysis with DESeq2 after excluding taxa with structural zeros; D. Analysis with DESeq2-ZINBWaVE after excluding taxa with structural zeros.

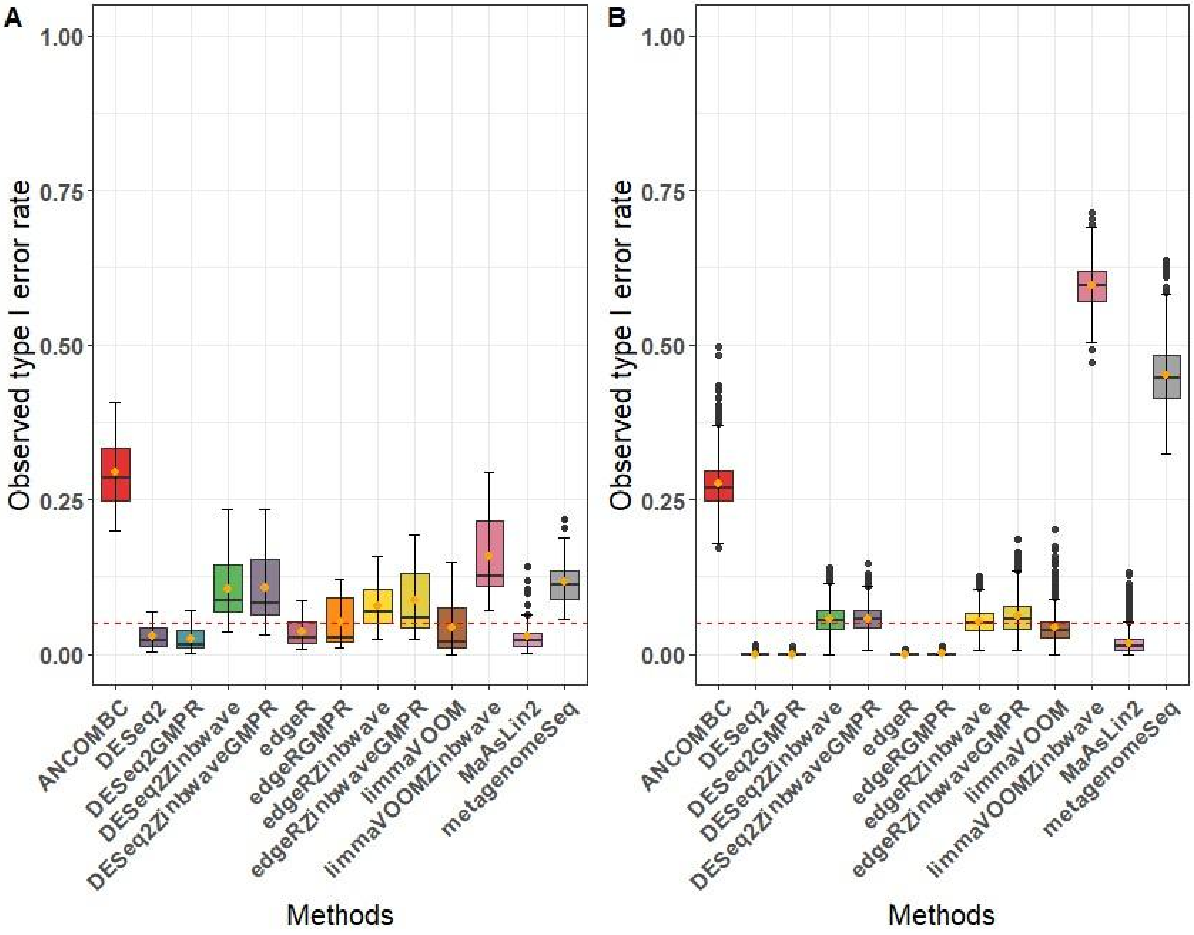


**Figure S2. Mock samples: controlling type I error rates using several differential abundance tools**. Unweighted and weighted differential abundance methods were evaluated for type I error control based on 1000 mock samples from plant microbiome data with varying zero-inflation rates. Left panel: N-P starvation template dataset with 55% zeros. Right panel: Forest-Potting soils template dataset with 84% zeros. Compared to the 5% nominal level, on average the observed type I error rates were very high for ANCOM-BC, metagenomeSeq, and limma-voom-ZINBWaVE; very low for DESeq2, edgeR and MaAsLin2; slightly higher for DESeq2-ZINBWaVE N-P starvation dataset;

and close to for DESeq2-ZINBWaVE Forest-Potting soils dataset and for edgeR-ZINBWaVE and under the null hypothesis of no differentially abundant taxa.

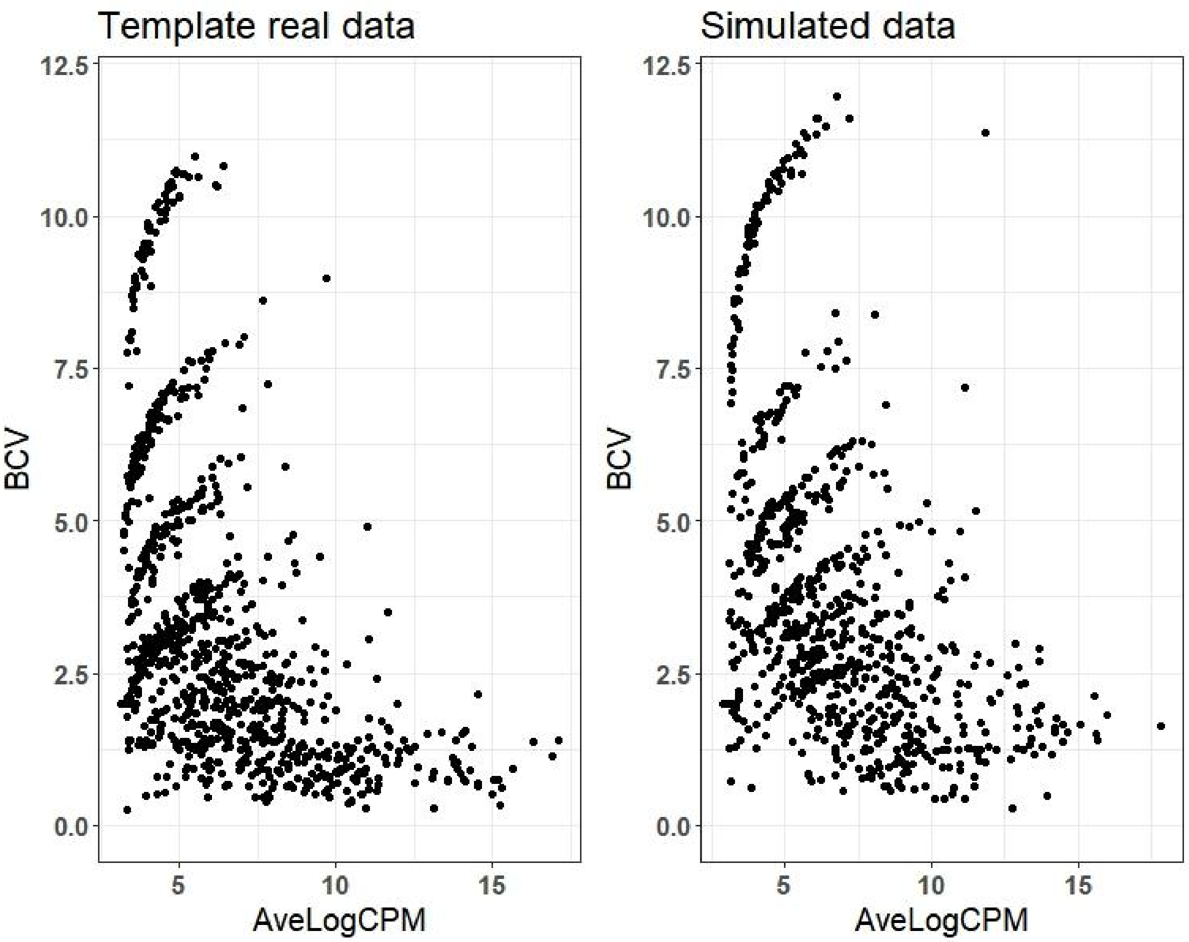


**Figure S3. Biological coefficient of variation (BCV) plots obtained from template real data and synthetic data**.The BCVs are displayed against the average number of log counts per million (AveLogCPM). The BCV plot displays striped patterns, which are a sign of taxa with many zeros and high estimates of dispersion. For the N-P starvation template real dataset, the BCV estimates from the template real data and one simulated data are comparable to each other.

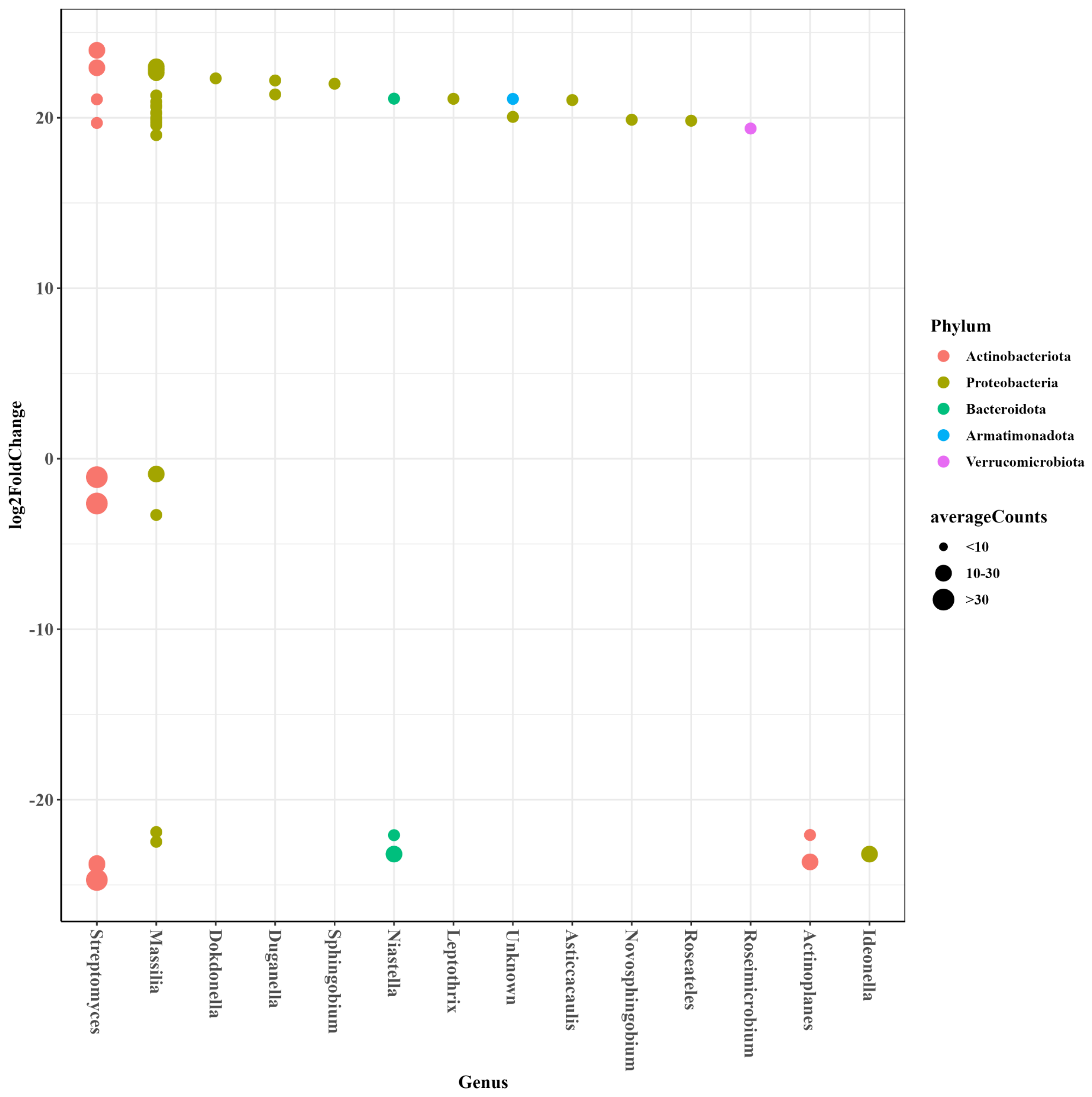


**Figure S4. Differentially abundant taxa in comparing potting and forest soils.** The size of dots indicate the average counts per taxa.

### Text S1. The potential of utilizing a combined DESeq2-ZINBWaVE and DESeq2 approach for differential abundance analysis

The simulation findings showed that, after removing taxa with perfect separation or group-wise structured zeros, which allowed for a fair comparison of methods, DESeq2-ZINBWaVE is found to be most powerful as a differential abundance analysis technique for substantially as well as moderately inflated microbiome data. This can be reinforced further by analyzing taxa with group-wise structured zeros using the DESeq2 likelihood ratio test, a tool that performs penalized likelihood inference and is well-suited for assessing taxa with group-wise structured zeros.

Therefore, as shown in Fig.1, we suggested to utilize a combined approach, which we refer to as DESeq2-ZINBWaVE-DESeq2 that includes a differential abundance analysis of taxa with group-wise structured zeros using the DESeq2 likelihood ratio test and taxa without group-wise structured zeros using the DESeq2-ZINBWaVE likelihood ratio test.

To assess the performance of the combined approach DESeq2-ZINBWaVE-DESeq2, we reanalyzed the metagenome shotgun sequencing data from the Human Microbiome Project (HMP-2012), which include 5 supragingival and 5 subgingival plaque samples from the oral cavity [1,2]. The data were utilized in enrichment analysis to rank methods according to how well they could detect taxa that are known to be differentially abundant between the supragingival and subgingival plaques [1]. After filtering with a minimum of 10 non-zero counts in at least 2 samples, 516 taxa were retained, of which 94 had group-wise structured zeros and 322 did not. Based on genus-level metabolism, each taxon was classed as aerobic, anaerobic, facultative anaerobic, or unclassified [1]. In comparing differential abundance of taxa between supragingival and subgingival plaques, it is expected to find an abundance of aerobic microbes in the supragingival plaque and of anaerobic microbes in the subgingival plaque. Fig. 6 shows the number of taxa found differentially abundant with adjusted p-value $< 0.10$ using DESeq2-ZINBWaVE-DESeq2 (134), edgeR-ZINBWaVE-edgeR (89), DESeq2-ZINBWaVE (70), and edgeR-ZINBWaVE (38). In Fig. 6, we displayed the number of significant taxa belonging to aerobic and anaerobic metabolism only. In this figure, the number of taxa with positive log-fold changes that are more abundant in the supragingival plaque are indicated by the top bars whereas the number of taxa with negative log-fold changes that are more abundant in the subgingival plaque indicated by the bottom bars. The color of the bars represent aerobic (red) and anaerobic (blue) metabolism. The top red bars demonstrated that all four approaches correctly identified aerobic taxa in the supragingival plaque. Moreover, all of the approaches correctly (bottom blue bars) identified anaerobic taxa with varying numbers in the subgingival plaque but they also incorrectly (top blue bars) identified a few anaerobic taxa in the supragingival plaque. Further, similar to Calgaro and colleagues [1], we performed enrichment analysis based on Fisher's exact test (see details in [1]) to explore the relationship between supragingival and subgingival plaques and the growth of aerobic and anaerobic microbial species. Using Fisher's exact test (p-values for significant enrichment indicated on the bars), we found that all of the methods considered here correctly found an enrichment of aerobic microbes among the taxa over-abundant in supragingival plaque (red bars) and an enrichment of anaerobic microbes among the taxa over-abundant in subgingival plaque (bottom blue bars). However, Fisher's exact test revealed no significant enrichment of anaerobic microbes among taxa over-abundant in supragingival plaque (top blue bars in Fig. 6). In comparing the four methods, the methods can be ranked according to the number of accurately identified taxa. As a result, the combined approaches accurately discovered many more anaerobic microbes enriched in the subgingival plaque (long bottom blue bars). In particular, we found DESeq2-ZINBWaVE-DESeq2 to be a more effective tool for discovering differentially abundant microbial species.

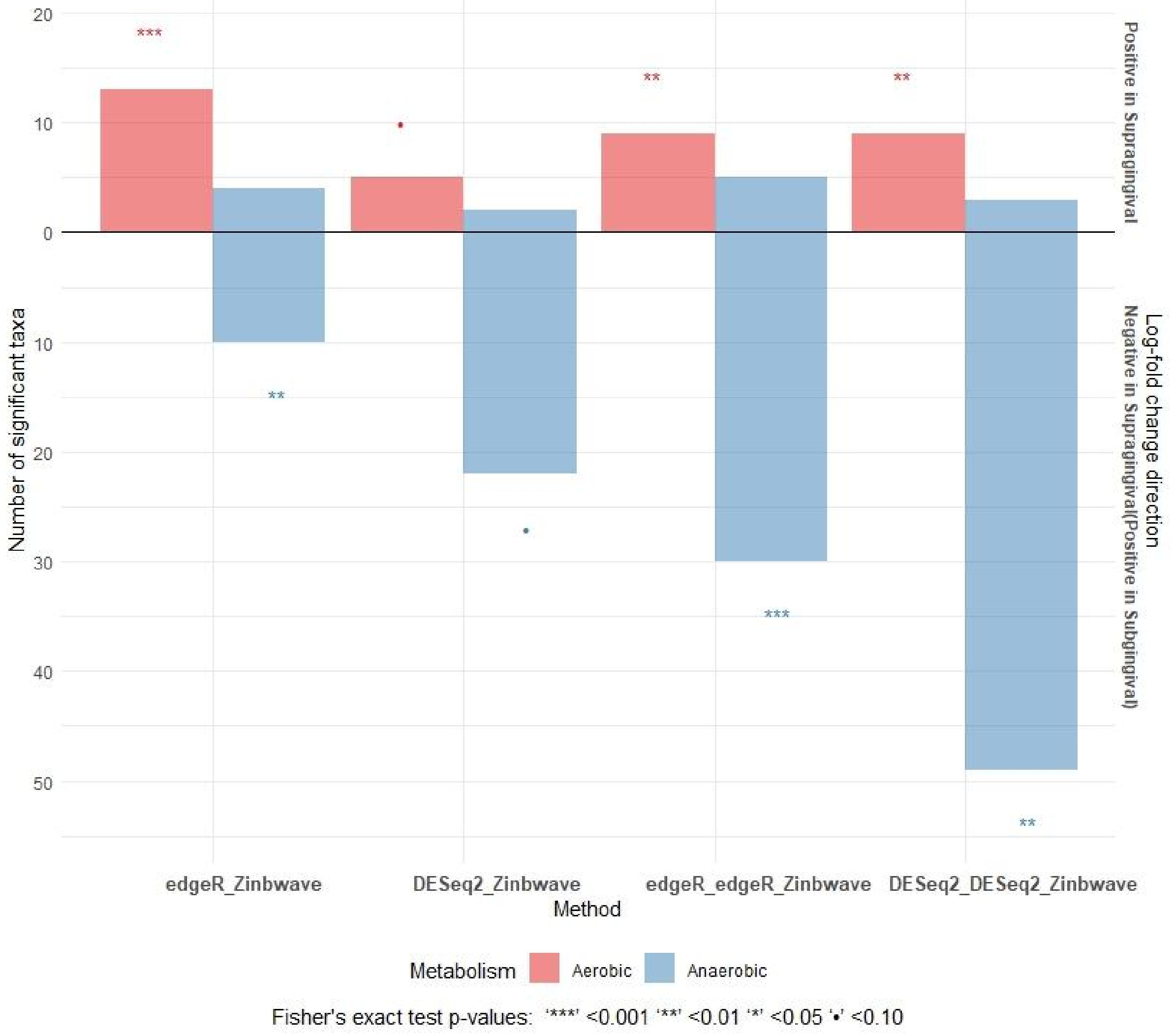


**Figure S5. Enrichment analysis comparing Supragingival vs Subgingival plaque for the metagenome shotgun sequencing samples from the Human Microbiome Project (HMP-2012).** Each bar represents the number of significantly (adjusted p-value < 0.10) abundant taxa by each method, with positive log fold changes in Supragingival (top red bars), negative log fold changes in Subgingival plaque (bottom blue bars), and positive log fold changes in Supragingival (top blue bars), coloured according to aerobic and anaerobic metabolism. A Fisher exact test is employed to establish the enrichment significance, and the p-values are shown.
